# Multi-omics Characterization of Duck Embryonic Stem Cells for Cultivated Meat

**DOI:** 10.64898/2026.04.27.720974

**Authors:** Remy Kusters, Théo Mathieu, Franks Kamgang Nzekoue, Matteo Manzati, Joao Palma, Brian Chun, Hannah Lester

**Author notes:** **Corresponding author:** Remy Kusters, PARIMA (SUPRÊME), 91058 Évry-Courcouronnes, France,.

## Abstract

Industrializing cultivated meat requires cell lines with high proliferative capacity, genetic stability, and suspension adaptability. We present a comprehensive multi-omics framework, integrating genomics, transcriptomics, and proteomics, to characterize a commercial duck Embryonic Stem Cell (dESC) line. Our analysis demonstrates continuous proliferation in protein-free suspension media while maintaining a stable genome and a functional conserved transcriptome. Broad-scale transcriptomics confirms the absence of hazardous pathway activation, and targeted assays verify sustained pluripotency marker expression during scale-up. Compositional analysis reveals a low-fat biomass containing all nine essential amino acids with an amino acid profile comparable to conventional duck meat. Furthermore, proteomic profiling demonstrates inter-batch reproducibility and protein distributions comparable to duck breast and liver. This study provides the first detailed molecular characterization of a commercial cultivated meat cell line, establishing a reference for the stability and safety assessment of future cultivated meat cell lines.

## Introduction

The industrial-scale production of cultivated meat presents significant biotechnological challenges, primarily centered on the development and characterization of robust animal cell lines. For cultivation to move from lab-scale proof of concept to large-scale manufacturing, cell lines must possess high intrinsic proliferative capacity and genetic stability while maintaining adaptability to suspension culture^1^. Ensuring the longitudinal consistency of these biological systems is critical for consistent bioprocessing and product safety. However, comprehensive data regarding the molecular profiles and compositional stability of commercial-scale cell lines remain limited in the current literature. Addressing these technical requirements is essential for establishing standardized frameworks for the production of cultivated biomass.

The transition from lab-scale proof of concept to industrial manufacturing has been hindered by fundamental biological and engineering bottlenecks^2^. Foremost among these is the identification of a cell source that combines constitutive self-renewal capacity with genetic stability, suspension adaptability, and the ability to proliferate in low-cost, food-grade media^3^. Muscle satellite cells have for a long time been the standard for cultivated meat due to their direct isolation and natural myogenic capacity^4^. To circumvent the replicative senescence of primary cells, the field has also relied on immortalized cell lines which employ genetic modifications to enable the indefinite proliferation required for industrial scaling^4^. These immortalized cell lines often possess unstable genomes and may require extensive genetic engineering to overcome senescence barriers, triggering more regulatory scrutiny and consumer aversion regarding Genetically Modified Organisms (GMOs) ^5^.

Embryonic Stem Cells (ESCs), and in particular avian ESCs, represent a safe and economically viable solution to these limitations. Unlike many mammalian counterparts, avian ESCs derived from the blastoderm stage of embryos exhibit a natural, intrinsic high proliferative capacity without the need for genetic engineering^6,7,8^. To demonstrate this, we provide a comprehensive multi-omics characterization of a commercial duck (*Anas platyrhynchos*) ESC line. We demonstrate that this cell line has the ability to grow in a protein-free and chemically defined suspension media, a critical milestone for ensuring both process safety and cost-competitiveness at scale.

This paper presents the first detailed examination of a commercial avian ESC line through a multi-omics framework covering genomics, transcriptomics, targeted pluripotency assays and compositional analysis. Multi-omics frameworks which integrate transcriptomic, metabolomic, and proteomic data have emerged as systems-level tools for mapping interactome networks and optimizing cultivated meat production^9,10^. These integrative approaches have been used to characterize the stability and metabolic flexibility of various cell sources, using models such as immortalized ovine satellite cells to evaluate the safety of novel production strategies^11^. Such high-resolution molecular characterization helps navigating the complex safety and regulatory requirements of the global cultivated meat industry ^5^.

We established the genomic integrity of the dESC line using Whole Genome Sequencing (WGS), to assess whether potential structural abnormalities or deleterious genetic drifts occurred across extended passaging. While WGS ensures a stable genetic blueprint, functional stability was validated through transcriptomic profiling. This analysis aims to confirm that key metabolic pathways remain consistent throughout the production process, showing negligible phenotypic drift. These studies were complemented by targeted RT-qPCR and flow cytometry, to verify the sustained expression of core pluripotency markers (e.g., *OCT4, NANOG*) throughout the cultivation lifecycle. Finally, we conduct a detailed proximate analysis to determine the macronutrient composition and evaluate the nutritional quality of the final biomass by profiling all nine essential amino acids. We employ a high-resolution bottom-up proteomics workflow to characterize the protein size distribution and inter-batch reproducibility, while performing a computational allergenicity assessment to verify that the production process does not introduce novel allergenic risks relative to conventional duck tissues.

## Results

### Establishment and Characterization of a Master Cell Bank

Industrial manufacturing robustness depends entirely on the quality of the starting material. To ensure long-term consistency, we established a tiered cell banking system (MCB and WCB) in adherence with ICH Q5D guidelines^12,13^. These banks underwent rigorous characterization to confirm identity and purity. Species identity (*Anas platyrhynchos*) was verified via PCR, ruling out cross-contamination with common biopharmaceutical cell lines^14^. To address regulatory safety requirements, we further screened the MCB using a validated real-time PCR panel targeting 38 viral pathogens, including avian-specific agents and potential bovine, porcine, or rodent contaminants. This targeted approach was complemented by agnostic, untargeted stranded RNA-seq on both the MCB and End-of-Process (EOP) samples. No evidence of known or novel replicative viral species was detected in either the bank or the final biomass, confirming the line is free of adventitious agents throughout the cultivation cycle. Trace alpharetroviral reads were identified but interpreted as endogenous retroviral transcripts expected under poly-A-selected duck RNA-seq.

The dESC line was derived and adapted to suspension culture through proprietary, non-GMO processes designed to select for high-fidelity stemness without genetic intervention. The process initially mimics the embryonic niche to support rapid proliferation before transitioning the cells to fully chemically defined, serum-free media^8,15,16^. This transition eliminates the safety and variability risks associated with animal-derived components like Fetal Bovine Serum. The resulting cells grow as non-adherent aggregates (**Figure 1A**), a morphology that protects against shear stress in large-scale bioreactors. Unlike many mammalian models, these avian cells maintain stable expansion and high pluripotency marker expression naturally due to distinct telomere maintenance mechanisms, precluding the need for exogenous DNA vectors ^8^.

**Figure 1.**
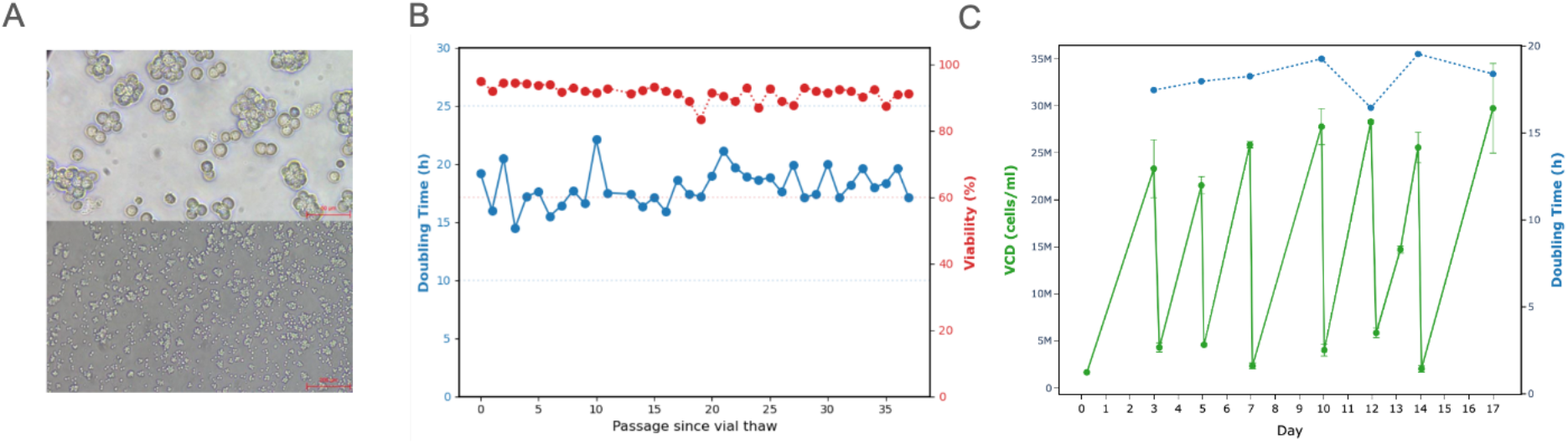
A) 40X and 20X magnification of the cell culture at passaging. Note that the top and bottom scale are 50 and 200 μm respectively. B) Doubling time and Viability post during the 90 days stability study. C) Viable Cell Density (VCD) and Doubling time as a function of time of cultivation in a semi-continuous process of 17 days.

### Phenotypical stability from vial to harvest

To demonstrate the phenotypical and functional stability of the dESCs we performed a 90-days stability study where the cells were continuously passaged in a shake flask, seeded at low Viable Cell Density (VCD: 0.3-0.5 Mcells/ml). In **Figure 1B** we show the doubling time and the viability of the cell culture for 37 passages (with a 2-3 days interval), indicating a stable Doubling Time (DT) of 18. 0 ± 1. 6 h and viability 91. 4 ± 1. 2% (See Methods). Cells were seeded at low cell density to avoid nutrient depletion and maintain a stable growth. We also verify that visually the cells maintain their round undifferentiated characteristics as shown in **Figure 1A**.

In a production setting, the expansion process of the cells can begin at any time within this window of established stability (90 days). Cell expansion consists of a series of expansion steps transferring the culture from shake flasks into progressively larger bioreactors. A scaled down production run was performed in a 10L bioreactor to mimic a semi-continuous process run and to assess the consistency of the doubling times and quantity of harvested biomass over multiple harvests. Throughout this expansion, phenotypical key performance indicators such as viability and visual inspection are tracked and are expected to remain consistent across scales and duration of the run. For this experiment, seven semi-continuous draw and fill cycles were performed with a 2-3 days interval. The seeding density here was increased compared to the 90 days study to improve yields (2.0-5.0 Mcells/ml). In **Figure 1C** we show the VCD and DT throughout this process. We find that the DT between seeding and harvest was 18.2 ± 1.1 h, consistent with the 90-days process in shake flask, where cells were seeded at much lower density. This demonstrates that even at commercially competitive cell performance, consistent yield is obtained. This highlights that no nutrient depletion occurs in the media throughout the culture. Visually the cells also remained identical throughout the process, no differentiation or cell death was observed. Taken together, all these phenotypical characteristics display a robust process stability.

### Whole Genome Sequencing

In order to support the assessment of genetic stability, Whole Genome Sequencing (WGS) was performed on the MCB and an EOP biomass sample. High-depth sequencing (∼30x coverage) was obtained for both samples (See Methods). This analysis confirmed that the global variant count relative to the reference genome (*Anas platyrhynchos*) remained consistent across all samples (∼7.6 million variants), indicating preservation of cell identity throughout the cultivation process.

The pairwise genetic divergence between samples was calculated at approximately 1 Single Nucleotide Polymorphism (SNP) per 3 kb. This variant density is consistent with the expected background heterogeneity of a diploid avian cell line derived from a domesticated lineage^17^. Crucially, this variant density remains stable between the MCB and EOP samples, confirming that the cell population maintains its genotypic identity without undergoing hyper-mutation accelerated drift during production.

To isolate mutations acquired strictly during the manufacturing process, a differential analysis of Private Variants (PVs) was performed. PVs are variants unique to a specific sample when compared to the reference sample. While the inherited genetic baseline remains consistent, the analysis identified only 639 to 666 high-impact PVs arising between the MCB and the EOP. This represents a negligible fraction (less than 0.01%) of the ∼7.6 million total variants in the genome.

Analysis of the Heterozygous-to-Homozygous (Het/Hom) SNP ratio reveals a highly consistent range of 1.73 to 1.82 across samples. This dominance of heterozygous variants confirms that the cell lines maintain a stable genome structure throughout the production process. A ratio of this magnitude is characteristic of stable outbred populations and indicates that the cell lines have not undergone significant Loss of Heterozygosity (LOH) or chromosomal instability^18^. The observed values reflect standard somatic drift and the accumulation of expected passenger mutations, rather than directed selective pressure. Because this biological system is engineered for terminal harvest for use in food, these discrete structural variations are consistent with expected background somatic variation over the tested production window and do not indicate progressive genomic instability, supporting the functional safety profile established in our transcriptomic assessments.

### Non-Targeted NGS Transcriptomics

While genotypic stability assesses the integrity of the genome itself, transcriptomic analysis (RNA-sequencing) provides a functional view of the biological state (transcriptome) of our dESC line. This approach enables a direct comparison between our cell line, developmental tissues, and mature cell lineages such as muscle and liver^19^. Moreover, assessing transcriptional consistency during the cultivation process and scale-up allows us to ensure that inadvertently harmful pathways are not activated ^5^.

To compare transcription across samples, RNA-sequencing was performed on a total of 113 samples. These included biological replicates of the WCB and in-process samples (at days 1, 4, and 7 of a batch cultivation). Additionally, we collected samples during early (0, 24 h, and 36 h after laying) and late (5, 10, and 13 days) duck embryo development (see Methods). We also incorporated RNA-seq data from different duck tissues available on the NCBI website (see Methods). Taken together, these diverse samples facilitate a comprehensive comparison between our dESC line and various reference tissues and development samples.

To evaluate the global gene expression profiles across these samples we selected the top 50% genes with most variance across samples. On this subset we performed a Principal Component Analysis (PCA) classifying the data into five groups: dESC line (yellow), early-stage embryo development (blue), late-stage embryo development (red), liver (green), and muscle (purple). As shown in **Figure 2A**, the two main principal components capture the majority of the variance, with PC1 accounting for 25.8% and PC2 for 23.2%.

**Figure 2.**
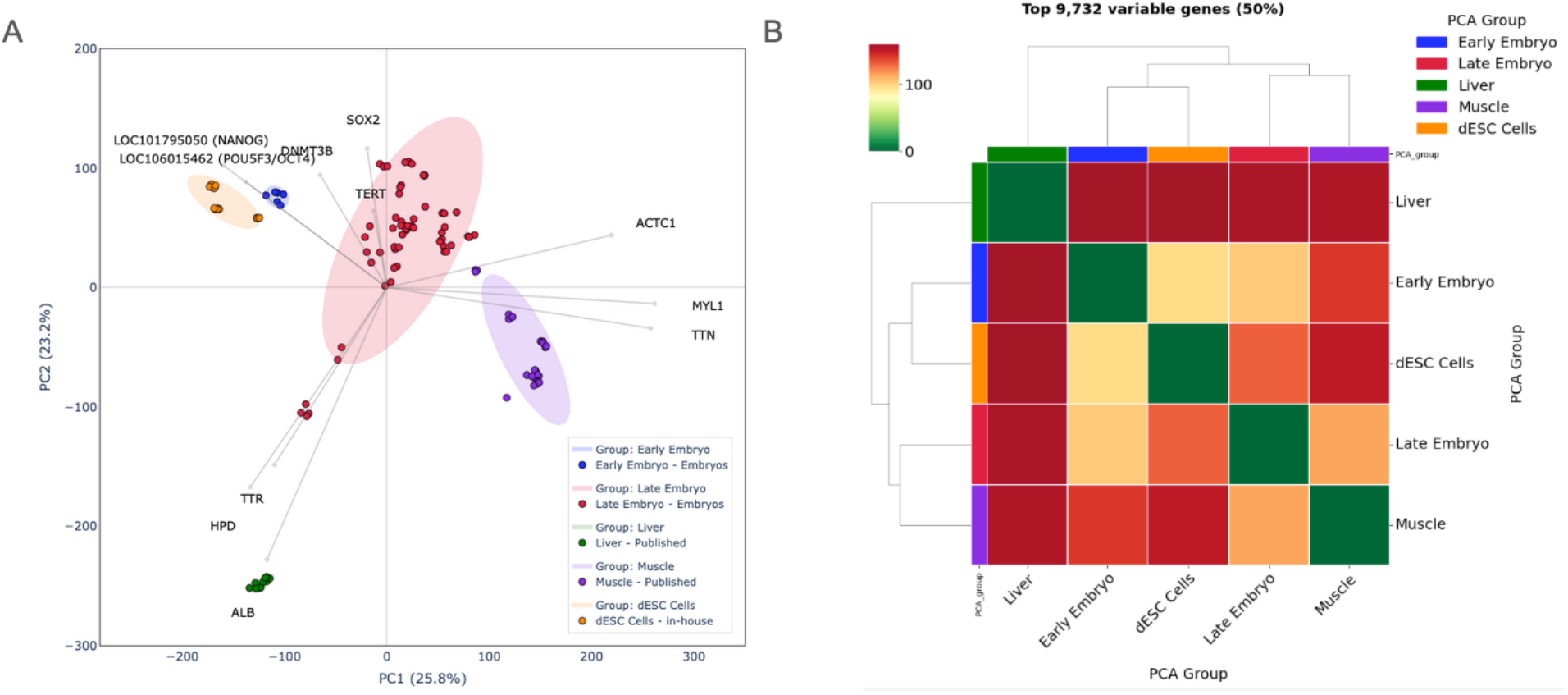
**A)** PCA of VST-normalized expression using the top 50% most variable genes. PC1 (25.8% variance) and PC2 (23.2% variance) show sample clustering by biological group (dESC Cells, Early/Late Embryo, Muscle, and Liver) with 2.0 SD confidence ellipses. Vector arrows indicate the influence of specific markers, including pluripotency factors (SOX2, NANOG, and POU5F3/OCT4), regulatory genes (TERT and DNMT3B), and differentiation markers (TTR, HPD, ALB, MYL1, TTN, and ACTC1). B) Euclidean distance mapping based on averaged group expression profiles. The analysis quantifies global transcriptomic divergence, highlighting the relative distance of embryonic stages and specialized tissues from the dESC Cells reference state.

To quantitatively assess the relative distance between these groups in the global expression landscape, we calculated the Euclidean distance across all conditions. This analysis demonstrates that the dESCs are most similar to early embryos, followed by late embryos and muscle, and are furthest from fully developed liver and muscle tissue (**Figure 2B**). This distinction is expected, as the cultivated biomass consists of undifferentiated ESCs rather than the highly structured matrix of differentiated muscle and connective tissues found in a mature organism^20^.

To further characterize the cells within this dimensionally reduced space, we quantified the expression of targeted biomarker genes. To contrast undifferentiated and differentiated states, we selected a panel of established biomarker genes. This panel includes core pluripotency maintenance genes such as NANOG (NCBI Gene ID: 101795050), POU5F3 (NCBI Gene ID: 106015462), TERT (NCBI Gene ID: 101791475), and DNMT3B (NCBI Gene ID: 101794583)^21^. NANOG is a transcriptional factor that suppresses cell differentiation^22^. POU5F3 induces cell proliferation^23^. TERT encodes for telomerase reverse transcriptase and is thus suppressed in differentiated cells and upregulated in stem cells^24^. DNMT3B encodes for a methyltransferase that is crucial in embryonic development and is downregulated during differentiation^25^. All four genes have been extensively used as markers of pluripotency^21,26,27^. We also included lineage-specific markers representative of distinct *in vivo* tissues, such as MYL1, ATCT1 and TTN for muscle and ALB, HPD and TTR for liver.

As indicated by the loading vectors (gray arrows in **Figure 2B**), the expression gradients for the pluripotency genes point uniformly toward the dESC and early-stage embryo clusters. Conversely, vectors representing differentiated tissue markers orient strictly toward the *in vivo* tissue and developed embryo references. This directional divergence visually confirms that the in-process ESCs retain a robust, pluripotent transcriptomic signature that is fundamentally distinct from the differentiated profiles of developed organs and late-stage embryos, consistent with the transcriptional dynamics observed during lineage commitment^28^. In the subsequent section, we will examine these pluripotency markers in greater depth using targeted assays.

Finally, we assessed the functional stability of gene expression during the cultivation process by performing a differential expression analysis between the WCB samples and the EOP samples (24 days of culture since vial thaw) obtained from a large-scale bioreactor. Comparing both transcriptomes, we identified transcripts corresponding to 284 genes that were more abundant in the end-of-process samples relative to the starter cells (WCB). While this shift likely reflects the cells’ adaptation from a post-thaw revival state from the WCB to a proliferation-adapted production state (EOP), as well as natural biological variability, the critical question is whether these transcriptional changes indicate the activation of hazardous biological functions. To address this, the upregulated genes were subjected to Gene Ontology (GO) enrichment analysis using GOATOOLS^29^. The analysis demonstrated that no biological pathways were statistically over-represented (FDR < 0.05).

The absence of statistically enriched GO terms (“Null Result”) is an indicator of functional stability. In biological systems, a functional shift, such as the activation of a toxin biosynthesis pathway or the onset of cellular stress, requires the coordinated upregulation of multiple genes within a specific pathway, rather than isolated gene variance^30^. A “Null Result” indicates that the 284 upregulated genes are stochastically distributed as transcriptional “noise” reflecting standard culture adaptation, rather than clustering into a directed biological “signal” of instability or hazard. Without pathway enrichment, individual upregulated genes lack the necessary enzymatic partners to drive a new metabolic phenotype. Specifically, there was no systematic activation of safety-relevant pathways, such as Oxidative Stress (GO:0006979) or DNA Damage Response (GO:0006974). These results complement our safety assessment, demonstrating that the production scale-up does not shift the cells toward a hazardous or unstable functional phenotype.

### Targeted Pluripotency Assessment

ESC-derived cell lines maintain the self-renewal ability, even when properly cultured *in vitro* for many passages^31,32^. Early studies showed that ESC-derived cells continue to proliferate to replace aging or damaged cells and do not differentiate into non-pluripotent cell types. For an industrial process where cell culture is progressively scaled-up, it is crucial to assess if this property is maintained throughout the process. In this section we assess pluripotency throughout the cultivation process through a dual approach that evaluates both the transcription levels of pluripotency-related genes via RT-qPCR and the expression levels of pluripotency-related protein markers via flow cytometry.

RT-qPCR serves as a method for measuring relative gene expression. The molecular markers identified to verify the pluripotent state of the culture are the same as those we used in the transcriptomics study: NANOG, POU5F3, TERT, and DNMT3B. The genetic expression of VIM, encoding for the structural protein vimentin associated with differentiated fibroblasts^33^, is used as a negative control. Finally, the accuracy of the assay was guaranteed by the use of two reference genes: RPS17 (NCBI Gene ID: 101798743), encoding for universal ribosome proteins, and GAPDH (NCBI Gene ID: 101803965), encoding for an enzyme crucial to glycolysis.

Targeted analysis demonstrated consistently high expression levels of core pluripotency markers relative to the housekeeping genes during continuous passaging (**Figure 3A**). The fold change in expression for key pluripotency markers in the cell line relative to a fibroblast negative control is coherent with the assumed embryonic cell type: NANOG displayed a 10^6^ fold change, POU5F3 (Oct4) a 10^4^ fold change, TERT a 10^2^-10^3^ fold change, and DNMT3B a 10^3^-10^4^ fold change. Conversely, the fibroblast-specific marker VIM was systematically low (10^-1^ fold change). Importantly, these expression levels were statistically equivalent between the stable phase of continuous passaging and the end of the production culture (“Passaging” and “Final vessel” in **Figure 3B** respectively). The fact that the level of expression of every gene remains stable indicates that the upregulation of pluripotency genes is maintained throughout the different stages of the expansion process, confirming that the manufacturing conditions do not induce significant changes in pluripotency or differentiation.

**Figure 3.**
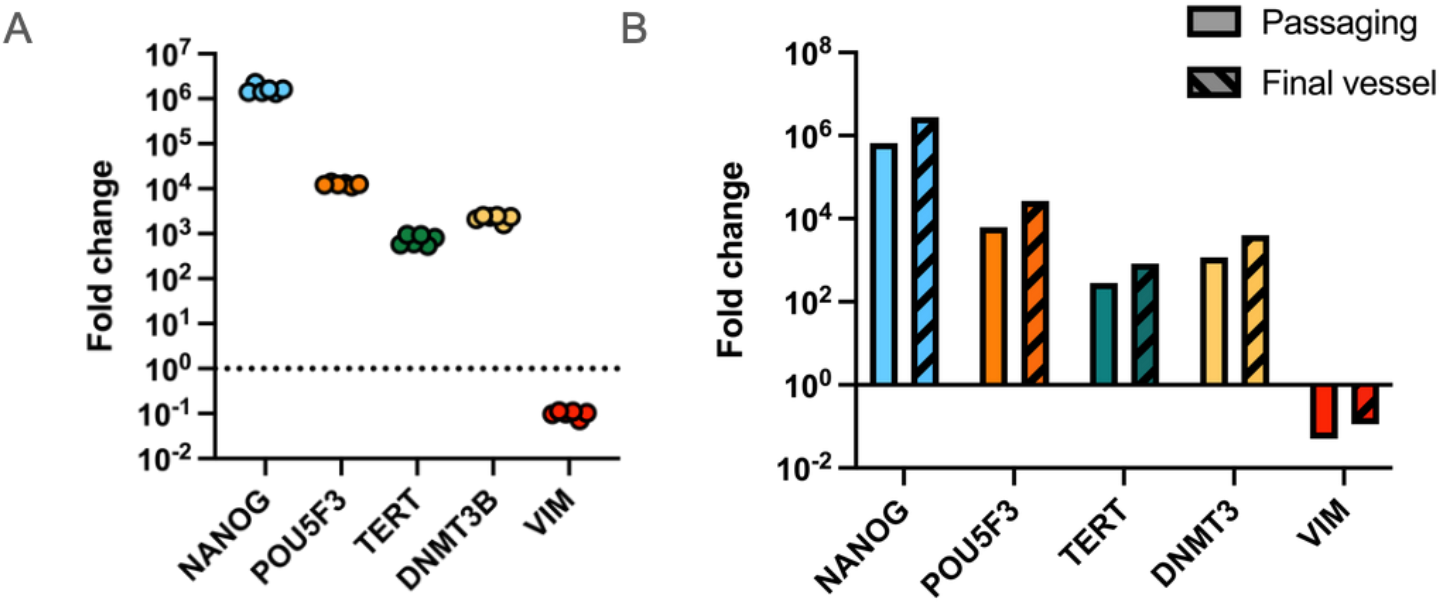
RT-qPCR. A) Fold change of gene expression for the genes NANOG, POU5F3, TERT, DNMT3, and VIM in the MCB (n=6). Cq values for each sample-gene combination has been normalized by the geometric mean of the Cq of two housekeeping genes (GAPDH and RSP17), then further normalized by the expression of the same gene in a negative control (CCL-114 duck fibroblasts). B) Fold change calculated as in A), but this time the same culture is compared at two different stages in the cultivation: Early stage shake flask passaging (Passaging) and prior to harvesting (Final vessel).

Flow cytometry provides validation that these genetic instructions are translated into functional intracellular and cell-surface proteins. It is a technique used to quantify the exact percentage of viable cells that express the markers of interest at the time of sampling. Two markers, EMA-1 and SSEA-1 have been reported to be highly relevant markers for avian ESC culture characterization^34,35^. The primary antibodies are subsequently detected by a secondary antibody conjugated to a fluorescent dye.

As shown in **Figure 4A**, the dESC cells, during continuous passaging, showed 38±5% positivity for EMA-1 and 48±8% positivity for SSEA-1, compared to <1% in negative control same-species fibroblasts (CCL-141). The assay demonstrated precision within the assay-accepted standard of CV < 25%^36^. A t-test analysis showed a significant difference between the dESCs and the negative control group (p = 0.0002 for EMA-1 and p = 0.0004 for SSEA-1).

**Figure 4.**
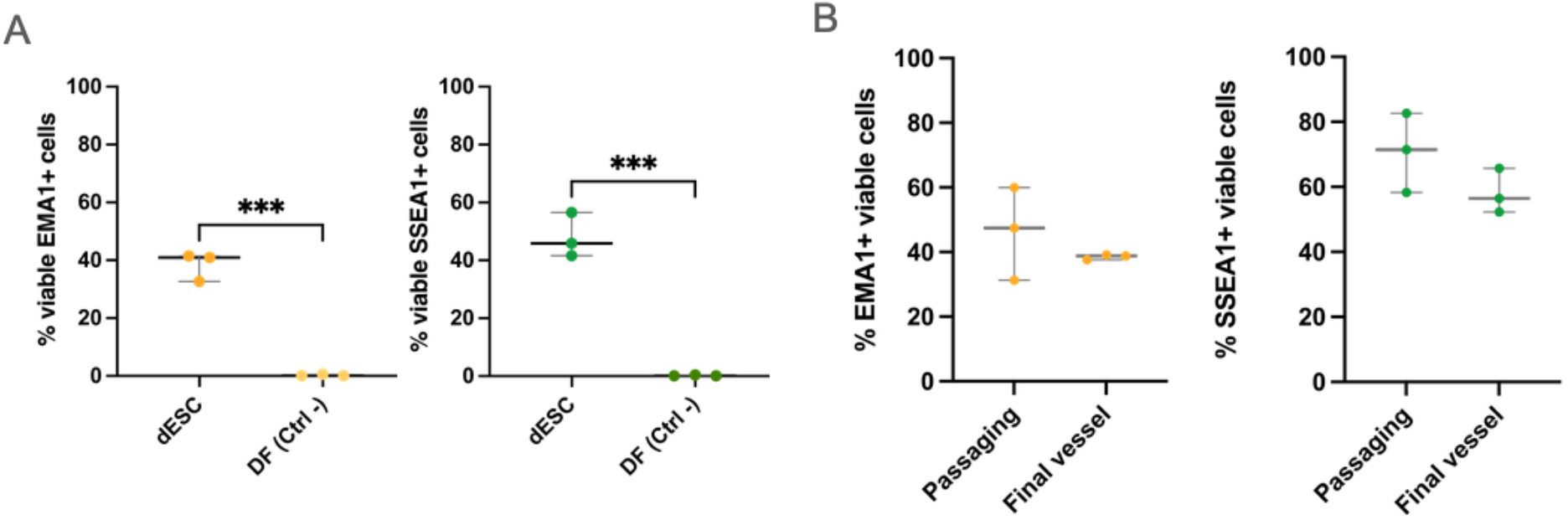
Flow Cytometry. A) % of detected duck embryonic stem cells (dESC) and CCL-114 duck fibroblasts (DF) that are both viable and EMA1+ (Left) and SSEA1+ (Right). Shapiro-Wilkis Normality test. Unpaired t-test, p = 0,0002 (EMA1+) and p = 0,0004 (SSEA1+) ; n = 3 technical replicates for each condition. B) % of detected duck cells that are both viable and EMA1+ (Left) and SSEA1+ (Right) as in A), but this time the same culture is compared at two different stages: Early stage shake flask passaging (Passaging) and prior to harvesting (Final vessel).

Positivity for both EMA-1 and SSEA-1 is also maintained when comparing continuous passaging (Passaging) and prior to final harvest of the cell culture (Final vessel), indicating that during the expansion process cells also maintain their intracellular and cell-surface pluripotency markers (**Figure 4B**).

The combination of the transcriptomics study with RT-qPCR and flow cytometry acts as a holistic confirmation of the pluripotent nature of the dESCs. RT-qPCR provides a sensitive, quantitative measure of the cell’s internal programming, while flow cytometry confirms the phenotypic “pluripotency” of the population at the protein level. The stability of these markers serves two purposes: i) it confirms the cells remain true ESCs throughout the process, and ii) it acts as a critical control for process drift.

### Proteins and amino acid profile

The nutritional quality of the final product is a core metric for evaluating cultivated meat. To ensure that avian ESC-derived biomass provides adequate protein quality we performed protein analysis from five production batches (*n*=5) of dESC biomass, comparing the results to conventional duck meat. Macronutrients analysis demonstrates that dESC biomass is nutritionally similar to conventional meat, being mostly composed of protein and moisture, and low in fat, consistent with its origin as undifferentiated ESCs. Notably, the average protein content across batches was approx. 12.0% (based on Kjeldahl (N x 6.25)) (**Table 1**). Furthermore, the amino-acid profile of ESC proteins confirms that the cultivated biomass contains all nine indispensable amino acids (IAA), which cannot be synthesized by the human body and must be obtained from the diet. Moreover, the overall amino acid (AA) profile remains consistent across batches, demonstrating the robustness of the production process. **Table 2** details the amino acid composition of cultivated dESC biomass and proteins in comparison to conventional duck. Although the total protein content is lower in dESCs (12.0g/100g) compared to conventional duck meat (18.5g/100g), the AA profile and the concentration of AA per gram of protein are comparable between the two. For instance, the order of abundance of individual AA remains consistent between duck and cultivated biomass. Glutamic acid is the most prevalent AA in both, followed by Aspartic acid and Lysine, whereas Tryptophan and Cysteine represent the least abundant AA across both sources. Furthermore, while conventional duck meat exhibits higher concentrations of specific amino acids — including Cysteine (281 vs 159 mg/100g), Phenylalanine (741 vs 457 mg/100g), Threonine (781 vs 471 mg/g), and Valine (956 vs 576 mg/100g) — the relative distribution per gram of protein remains highly comparable across both sources. Comparative analysis between conventional duck and cultivated dESC biomass shows that values for Cysteine (15.4 vs 13.3 ± 4.4 mg/g protein), Phenylalanine (41.9 vs 38.1 ± 7.8 mg/g protein), Threonine (42.7 vs 39.2 ± 8.4 mg/g protein), and Valine (52.3 vs 48.0 ± 10.1 mg/g protein) fall within comparable ranges.

**Table 1.** Proximate compositions of cultivated duck ESC biomass in comparison to other meat products.

| Compositional parameters | Units | dESC biomass (mean ± SD)<sup>1</sup> | Duck meat<sup>2</sup> | Goose liver<sup>2</sup> |
| --- | --- | --- | --- | --- |
| Energy | kcal/100 g | 57.4 ± 13.8 | 135.0 | 133.0 |
| Water | g/100g | 85.2 ± 1.7 | 73.8 | 71.8 |
| Protein | g/100g | 12.0 ± 1.2 | 18.3 | 16.4 |
| Total Lipid (fat) | g/100g | 1.2 ± 0.5 | 6.0 | 4.3 |
| Saturated fats | g/100g | 0.7 ± 0.0 | 2.3 | 1.6 |
| Total Carbohydrate | g/100g | < 0.1 | 0.0 | 6.3 |
| Ash | g/100g | 1.4 ± 0.3 | 1.1 | 1.3 |

<sup>1</sup>Values of dESCs biomass obtained from five culture batches ( $n = 5$ ).
<sup>2</sup>Values retrieved from FoodData Central (U.S. Department of Agriculture – USDA).

**Table 2.** Amino Acid Profile of Cultivated Duck ESC biomass in comparison to conventional meat (duck breast and goose liver). Values for cultivated meat are expressed as mean ± standard deviation (*n*=5)

|  | Amino acid concentrations<br>(mg/100 g product) |  |  | Amino acid concentrations<br>(mg/g protein) |  |  |
| --- | --- | --- | --- | --- | --- | --- |
|  | Cultivated<br>dESC<br>biomass | Duck Meat <sup>1</sup> | Goose Liver<sup>1</sup> | Cultivated dESC<br>biomass | Duck Meat<sup>1</sup> | Goose Liver<sup>1</sup> |
| **Indispensable Amino Acids (IAA)** |  |  |  |  |  |  |
| Histidine | 256.2 ± 52.2 | 483 | 435.0 | 21.4 ± 4.4 | 26.4 | 26.5 |
| Isoleucine | 472.8 ± 99.6 | 939 | 870.0 | 39.4 ± 8.3 | 51.4 | 53.0 |
| Leucine | 879.2 ± 185.1 | 1544 | 1480.0 | 73.3 ± 15.4 | 84.5 | 90.2 |
| Lysine | 906.4 ± 165.8 | 1564 | 1240.0 | 75.5 ± 13.8 | 85.6 | 75.6 |
| Methionine | 240.0 ± 55.7 | 494 | 388.0 | 20.0 ± 4.6 | 27.0 | 23.7 |
| Phenylalanine | 457.4 ± 93.9 | 766 | 815.0 | 38.1 ± 7.8 | 41.9 | 49.7 |
| Threonine | 470.8 ± 100.4 | 781 | 728.0 | 39.2 ± 8.4 | 42.7 | 44.4 |
| Tryptophan | 143.7 ± 28.1 | 254 | 230.0 | 12.0 ± 2.3 | 13.9 | 14.0 |
| Valine | 576.0 ± 121.7 | 956 | 1030.0 | 48.0 ± 10.1 | 52.3 | 62.8 |
| **Non-Indispensable Amino Acids (Non-IAA)** |  |  |  |  |  |  |
| Cysteine<sup>2</sup> | 159.0 ± 52.6 | 281 | 220.0 | 13.3 ± 4.4 | 15.4 | 13.4 |
| Glycine | 578.0 ± 33.1 | 1024 | 951.0 | 44.3 ± 9.0 | 56.0 | 58.0 |
| Aspartic acid | 1107.5 ± 29.9 | 1790 | 1560.0 | 84.8 ± 16.9 | 97.9 | 95.1 |
| Glutamic acid | 1532.5 ± 79.3 | 2860 | 2120.0 | 117.4 ± 23.7 | 156.5 | 129.3 |
| Proline | 517.5 ± 18.9 | 898 | 812.0 | 39.5 ± 8.3 | 49.1 | 49.5 |
| Serine | 533.0 ± 22.5 | 787 | 705.0 | 40.7 ± 8.5 | 43.1 | 43.0 |
| Tyrosine | 391.0 ± 77.0 | 594.0 | 696 | 32.6 ± 6.4 | 38.1 | 35.1 |
| Alanine | 652.5 ± 31.9 | 1158 | 951.0 | 49.9 ± 10.2 | 63.3 | 58.0 |
| Arginine | 831.0 ± 38.7 | 1166 | 1000.0 | 63.6 ± 13.0 | 63.8 | 61.0 |

<sup>1</sup>Values retrieved from FoodData Central (U.S. Department of Agriculture – USDA).
<sup>2</sup>Values of Cysteine result from the sum of cysteine and cystine.

### Proteomics

Beyond the standard protein analysis, a proteomics study was conducted to characterize the protein profile of dESC biomass relative to duck breast and liver. For this purpose, three production batches of dESCs were collected and analyzed alongside duck breast and liver samples using a bottom-up proteomics workflow^38^. Analysis of the dESC proteome revealed exceptional stability across production batches, with less than 3% of variation in protein composition between the three study groups (**Figure 5A**). Principal Component Analysis (PCA) further validates this consistency, showing that the batches cluster tightly together and significantly differ from the breast and liver tissues (**Figure 5B**). This high level of reproducibility underscores the robustness of the dESC line and the optimization of bioprocess conditions for standardized biomass manufacturing.

**Figure 5.**
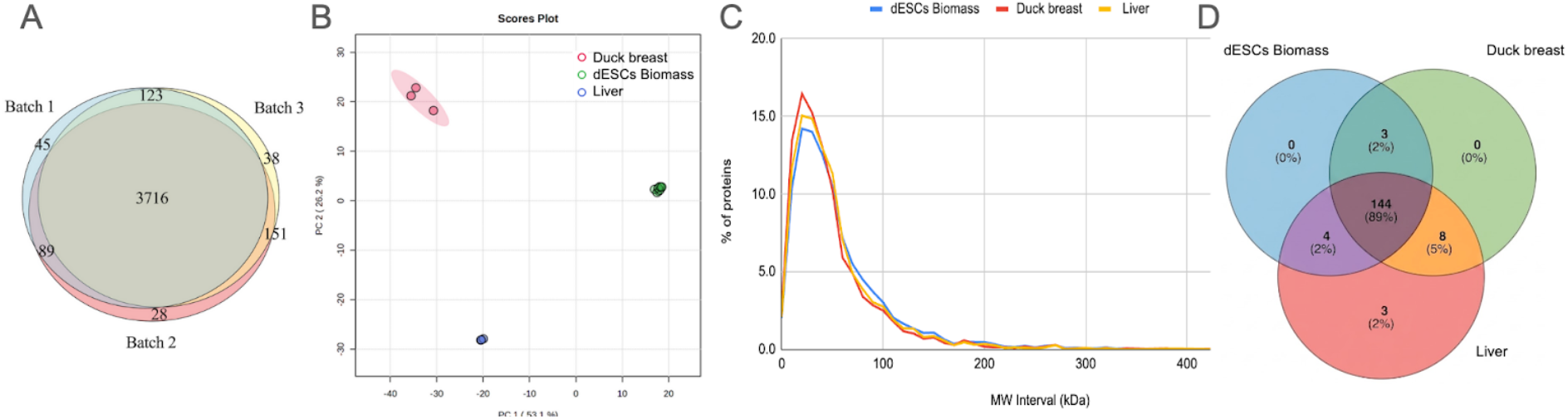
A) Venn diagram of protein groups identification across different batches of dESCs biomass; B) Principal Component Analysis (PCA) of protein-normalized abundances. PC1 (53.1% variance) and PC2 (26.2% variance) show sample clustering by biological group (dESC biomass, Duck breast, and Liver) C) Protein size distribution (MW) of dESCs biomass, duck breast, and liver proteins D) Venn diagram of matched allergens (subjectsID) against protein groups (QueryID) identified across different dESCs biomass, duck breast and liver. Alignment was performed through BLASTP alignment against a comprehensive allergen database (UniprotKB allergens_SP).

**Figure 5C** shows the protein size distribution of dESCs biomass, duck breast and liver in terms of Molecular Weight (MW) across sample groups. Globally, the protein size profile of the dESCs biomass are highly comparable to duck breast and liver tissues in terms of MW and Amino Acid (AA) length distributions (**Figure S1**). This protein size profile is characterized by a dominant presence of ∼10-50 kDa and 100-400 AA long proteins (∼70% of total proteins) followed by a rapid decline in frequency as protein size increases (60 - 200 kDa and 500 - 2000 AA). Duck breast tends to have a higher proportion of low MW proteins followed by liver and dESCs biomass.

While overall all samples have a comparable size distribution, differences in the proteome were observed. In terms of Proteins IDs, around 4190 protein groups were identified in dESCs vs 2933 in duck breast and 3951 in duck liver. This difference is likely attributable to the functional specialization of adult tissues, which result in higher abundance of fewer proteins with possibly lower apparent diversity due to the analytical dynamic range. Conversely, stem cells maintain a broader proteome as they express developmental regulators and multiple lineage priming pathways^39^. For example, in muscle tissue, actin and myosin can constitute between 40 – 60% of the total protein mass resulting in a lower detection of lower-expression proteins^40^. Specifically, duck breast, which is rich in muscle, resulted significantly high in Myoglobin and in proteins involved in cytoskeletal activity (e.g. Myosin heavy chain,Troponin I, Myotilin,, MYPN, and Tppp3) and kinase activity (e.g. Creatine kinase, pyruvate kinase, MYLK2). Duck liver was characterized by higher enzyme expression mainly transferases (e.g., GSTs, GGACT), oxido-reductases (e.g. GMP reductase), and Cytochrome P450 ^41^. dESC biomass showed a distinct enrichment in proteins associated with GO terms characteristic of actively dividing and undifferentiated stem cells – nucleus, nucleic acid binding, DNA/RNA metabolism, and the cell cycle (**Figure S2**). Moreover, dESCs biomass revealed a significant expression level of proteins linked to cytoskeletal activity including Actin-related proteins, Tubulins (e.g. alpha-chains, beta-chains), Tropomyosins (e.g. alpha-1, 1 – 4), and Myosins (e.g. 10, IXb, Va,).

### Allergenicity assessment

A computational allergenicity assessment was conducted to ensure that the production of cultivated meat does not introduce novel allergens absent in traditional duck meat^42^. From the proteomic results, amino acid sequence homologies of dESC biomass/ duck liver/ duck breast proteins against a comprehensive database of known allergens was determined (UniprotKB allergens_SP). Under this evaluative framework, proteins exhibiting more than 35% homology over a sequence of 80 or more amino acids with E-value of BLASTP search < 0.001, are considered to have a significant allergic cross-reactivity potential ^43^.

This analysis identified 162 matches to known allergens in the duck comparator tissues (≥ 35% homology over ≥ 80 AA), whereas the dESC biomass did not show additional sequence-homology alerts in the applied screen (**Figure 5D**). Proteins corresponding to known poultry allergen matches, such as ovotransferrin (sp|P02789|TRFE_CHICK), ovoinhibitor (sp|P10184|IOV7_CHICK), or chicken albumin (sp|P19121|ALBU_CHICK), were lower in dESC biomass than in duck tissues. In the applied sequence-homology screen, dESC biomass did not show additional allergenicity signals relative to the duck comparator tissues.

## Discussion and Conclusions

In this study, we have presented the first of its kind comprehensive multi-omics characterization of a commercial cell line for cultivated meat production, a duck Embryonic Stem Cell (dESC) line. We demonstrated that these cells provide a unique solution to the dual challenges of manufacturing efficiency and safety. By integrating genomics, transcriptomics, proteomics, and targeted analysis, we have constructed a multi-omics profile that moves beyond simple phenotypic observations and provides a high-resolution map of cellular behavior across the production lifecycle.

Our findings establish a new benchmark for cell line characterization. Whole Genome Sequencing (WGS) confirms that dESCs maintain a stable, diploid genome and consistent variant density from the Master Cell Bank through to the end-of-process harvest. This genomic stability is functionally confirmed in our transcriptomic landscape. Complementing these findings, our multi-tiered screening for adventitious agents, including a targeted PCR panel for 38 viral pathogens and untargeted stranded RNA-seq, on both WCB and EOP samples confirmed that no exogenous human or foodborne viral pathogens were detected in the analysed WCB and EOP samples.

The use of broad-scale RNA-sequencing and Principal Component Analysis (PCA) allowed us to verify that our cells retain a clear pluripotent identity, distinct from differentiated tissues, yet highly consistent across long-term cultivation. Crucially, the absence of statistically enriched GO terms among genes more abundant at the end of the process supports functional stability during the tested scale-up comparison, demonstrating that the pressures of scale-up do not induce deleterious functional shifts. The integration of targeted assays, RT-qPCR and flow cytometry, further strengthen this assessment by bridging the gap between genetic instruction and protein expression. By confirming that core pluripotency markers (NANOG, POU5F3, TERT, DNMT3B) and surface antigens (EMA-1, SSEA-1) remain stable throughout the cultivation process, we provide a redundant, high-fidelity method for identity verification.

The assessment of the amino acid profile and protein levels validates the strategy of dESCs as novel meat alternatives. Proteomics completed the multi-omics characterization of ESCs, providing a comprehensive understanding of the proteome, including size distribution, protein abundances, and biological relevance^44^. Taken together, those results emphasized the nutritional quality of dESC proteins with an allergenicity profile similar to conventional duck.

Beyond demonstrating the technical attributes of dESCs this study offers a contribution to the advancement of regulatory science. For risk assessors and regulatory agencies, the multi-omics framework presented here may serve as a foundation to inform future safety and risk assessment frameworks for cultivated foods. As the industry moves toward more complex production systems, integrated omics are increasingly important to demonstrate functional stability of the process and the safety and nutritional consistency of the end product^45,46^. By providing a transparent, molecular-level assessment of the cell line, we can shift the regulatory conversation from novelty to rigorous characterization, fostering the public trust necessary for the widespread adoption of cellular agriculture. This framework closely aligns with the New Approach Methodologies (NAMs), which emphasize human-biology-based, non-animal testing strategies to ensure product safety and quality ^47,48^. Such integrative models are increasingly advocated by global regulatory agencies, including the FDA and EFSA, to support tiered, weight-of-evidence safety assessments that prioritize molecular fidelity and compositional transparency over traditional *in vivo* toxicological studies^5,43,49^.

In conclusion, the avian ESCs represent a robust benchmark for the industry: a naturally immortal, non-GMO cell source that is metabolically optimized for suspension culture. The multi-omics framework presented in this paper supports characterisation of the tested dESC line across the sampled production window and may inform broader stability and safety assessments for the next generation of cultivated biomass, ensuring they are safe and nutritious.

## Methods

### Phenotypical Stability from cell to harvest

We assessed the cell growth by measuring the cell density, viability using the Norma XS (cell counting equipment) and calculating the doubling time *DT* = Δ*t ln*(2)/ *ln* (*c*_2_ /*c*_1_), where Δ*t* is the time interval between two passages in hours and *c*_2_ and *c*_1_ are the end and starting viable cell density respectively. The cell phenotype is assessed by measuring the cell diameter and visually inspecting the cell morphologies.

### Whole genome sequencing

#### Nucleic Acid Extraction and Library Preparation

Total genomic DNA, inclusive of both chromosomal and extra-chromosomal (mitochondrial) elements, was extracted from the cell pellets by an external party. The isolation of the high molecular weight (HMW) DNA was performed with the Wizard HMW DNA extraction kit. Genomic DNA concentrations were determined using a fluorescence-based assay, and quality was assessed using the QIAxcel Connect system prior to library construction. Sequencing was conducted on the Illumina NovaSeq X Plus sequencer using paired-end chemistry (2x150bp) to ensure high-resolution coverage of the genome. Base-calling was performed directly on the instrument using Illumina Real-Time Analysis (RTA) software Library construction was performed using the NEBNext® Ultra™ II FS DNA PCR-free Library Prep Kit for Illumina (cat# NEB#E7430S/L). Genomic DNA was fragmented via enzymatic fragmentation, followed by end repair, A-tailing, and ligation of sequencing adapters according to the manufacturer’s instruction manual.

#### Sequencing Strategy and Instrumentation

Clustering and DNA sequencing using the NovaSeqX plus was performed according to the manufacturer’s protocols. A final loading concentration of ∼160 pM of DNA was used. Sequencing was conducted using paired-end chemistry (151-16-9-151) to ensure high-resolution coverage of the genome. Base calling and initial quality checks were performed with the Illumina BclConvert v4.3.6.

#### Quality Control, Filtering Parameters, and Coverage Rationale

Raw sequencing reads underwent rigorous quality control and filtering using DRAGEN QC 4.0.6-1. Quality control metrics confirmed excellent sequencing performance, achieving a minimum per-base Phred score of 36.16, exceeding the standard acceptable threshold of Q20 for short-read sequencing.

The sequencing run achieved an average read depth of ∼25x to 31x across the ∼1.3 Gb *Anas platyrhynchos* genome. While the EFSA WGS statement for microorganisms recommends 30x to 100x depth for small microbial genomes, sequencing depth requirements scale differently for complex eukaryotes. According to historical benchmarks established for eukaryotic genomes, an average depth of just 15x is sufficient to detect almost all homozygous single-nucleotide variants, while a depth of ∼30x emerged as the de facto standard for comprehensive heterozygous variant detection ^50^.

Applying this benchmark to the current study, an average depth of ∼25x to 30x provides a high-confidence representation of both heterozygous and homozygous positions. This is supported by the fact that the *Anas platyrhynchos genome* (∼1.3 Gb) is substantially smaller than the human and mammalian models (∼3.0 Gb) used to establish historical 30x standards. Furthermore, the transition to patterned flow cell technology and modern chemistry utilized in this study improves the uniformity of read distribution across the genome. This reduction in coverage variance ensures robust statistical power for variant calling without requiring the excessive over-sequencing typically designed for *de novo* microbial assemblies.

#### Data Analysis

Following sequencing, we performed preprocessing, variant discovery, and quantification. The data analysis pipeline was executed as follows:

- **Quality Control:** Adapter and polyG sequences were removed via hard trimming.
- **Alignment:** Cleaned reads were mapped to the reference genome GCF_047663525.1_IASCAAS_PekinDuck_T2T_genomic using the DRAGEN platform.
- **SNPs and Indels:** Variant calling and genotyping were performed using DRAGEN. Metrics including the total number of SNPs, indels, transitions, transversions.
- **Hard Filtering:** To ensure high confidence, variant files were filtered following default parameters.

### Broad Scale NGS Transcriptomics

RNA-sequencing was performed on a total of 113 samples. These included biological replicates of the WCB and in-process samples (at days 1, 4, and 7 of cultivation). Additionally, we collected samples during early (0, 24 h, and 36 h after laying) and late (5, 10, and 13 days) duck embryo development. Below we will describe the procedures for RNA Extraction, Library Preparation, Sequencing as well as bioinformatic processing and functional analysis.

#### dESC samples

Total RNA was extracted from cultured dESC cells using either the Qiagen RNeasy Midi Kit (incorporating an RNase-Free DNase step) or the NucleoSpin RNA Mini Kit (Macherey-Nagel), depending on the experimental phase. Sequencing libraries were prepared using the Illumina Stranded mRNA Prep (poly-A selection), with an additional 10 cycles of amplification applied where necessary. All libraries were sequenced on an Illumina platform (NovaSeq) using 150 bp paired-end reads. Quality control via FastQC confirmed consistently high data quality across all samples, with >92.9% of bases exceeding Q30 and robust depths of >100 million reads per sample, ensuring high sensitivity for detecting low-abundance transcripts.

Since the datasets were generated during two independent experimental phases, they were processed using the standardized nf-core/rnaseq pipeline (v3.14.0). Reads were aligned to the Anas platyrhynchos reference genome assemblies (GCF_015476345.1_ZJU1.0 and GCF_047663525.1_IASCAAS_PekinDuck_T2T) using STAR (v2.7.11b) under default parameters. Gene-level quantification was conducted using Salmon via the star_salmon mode to estimate transcript abundance. Functional analysis was performed by grouping genes into molecular functions based on Gene Ontology (GO) classifications. Statistical enrichment of GO terms was calculated using GOATOOLS ^29^.

#### Early and late embryo samples

Total RNA was isolated using stage-specific protocols optimized for embryonic tissue complexity. For early-stage embryos (H0, H24, and H36), RNA was extracted using a modified Trizol/Chloroform method. Samples were homogenized in 400 µl of Trizol reagent (Thermo Fisher), followed by phase separation with chloroform and an additional chloroform wash to ensure high purity. RNA was precipitated using isopropanol with the addition of 1 µl of glycogen (Thermo Fisher) as a carrier, followed by sequential washes in 75% and 100% ice-cold ethanol. The resulting pellets were dried at 37°C for 5 minutes and resuspended in 15 µl of nuclease-free water. For late-stage embryos (D5, D10, and D13), tissues were homogenized in Trizol (1 ml per 75 mg tissue) and subjected to chloroform phase separation. The resulting aqueous phase was processed using the NucleoSpin RNA Plus kit (Macherey-Nagel), where RNA was bound to columns after the addition of 70% ethanol. Following optimized wash steps and membrane drying, RNA was eluted in 20 µl of nuclease-free water. For all samples, RNA concentration and purity were initially assessed using a NanoDrop One Spectrophotometer. RNA integrity was further validated using the Qubit RNA IQ Assay kit (Thermo Fisher) to ensure suitability for high-depth sequencing.

The bioinformatic analysis was performed by an external party using a robust pipeline for high-throughput transcriptomic data. A total of 6,957,177,170 paired-end reads (150 bp) from 63 Illumina libraries were generated for *Anas platyrhynchos*. Pre-processing was conducted using FastP for adapter removal and quality trimming, with FastQC employed to verify the quality of the cleaned reads. This stage demonstrated high data integrity, with a read survival rate ranging from 98.4% to 99.8%, resulting in 6,908,436,904 pairs of reads retained for downstream analysis. The cleaned reads were subsequently mapped to the *Anas platyrhynchos* reference genome (GCF_015476345.1) using STAR in paired-end mode. Mapping quality control was rigorously monitored via RNA-SeQC2, with mapping rates generally between 84% and 92%. Finally, transcript abundance estimation was achieved using the RSEM method to provide a high-resolution quantitative dataset for further investigation.

Muscle and liver tissue samples: The data can be retrieved on BioProject with below accession number and SRA identifier in the NCBI BioProject database:

PRJNA726590: SRR14398903–08

PRJNA878639: SRR21494906, SRR21494912

PRJNA349262: SRR4434780–97

PRJNA437527: SRR6820457–65

#### Data processing

Raw reads from all sources were concatenated into a unified dataset. To ensure robust downstream analysis and reduce technical noise, a strict filtering step was applied to retain only genes present in all of the component studies. The resulting count matrix (113 samples and 19465 genes) was then normalized to account for library size and heteroscedasticity. Normalization was performed using the Variance Stabilizing Transformation (VST) via the PyDESeq2 framework. The VST was calculated using a parametric fit type, which transforms the count data to a log-like scale where the variance is approximately independent of the mean. This approach provides a high-quality, homoscedastic dataset suitable for visualization and machine learning applications. The final VST-transformed counts were exported for further differential expression analysis.

### Targeted pluripotency assessment

*RT-qPCR*. Frozen pellets are thawed on ice. RNA extraction is performed using the commercially-available NucleoSpin RNA, Mini kit for RNA purification from Macherey Nagel (Reference 740955.5) kit. The extracted RNA is then converted into cDNA using the Maxima H Minus First Strand cDNA Synthesis kit according to company specifications. A SYBR Select Master Mix (Thermo Fisher Scientific) is then prepared, adding to it: reverse and forward primers for the following gene sequences: NANOG, POU5F3, TERT, DNMT3B, VIM and GADPH. The qPCR reaction mix and the cDNA at a final concentration of 2.5 ng/µL are loaded in a 384 wells plate. The plate is analysed with the QuantStudio 384: an activation step followed by 40 cycles of denaturation, annealing and elongation.

The quantification cycle (Cq), i.e. the number of cycles at which the sample’s reaction crosses the fluorescence threshold for SYBR green, is calculated for all markers for each sample. The Cq values are then normalized in ΔCq values:

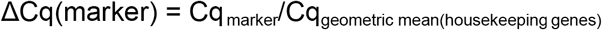

and the relative expression (RE) is calculated:

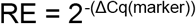

Finally, the fold change relative to a negative control, the commercially available duck fibroblasts (DF) cell line CCL-114 is calculated for all markers for each sample:

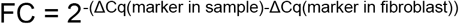

The results are then expressed as this Fold change (See **Figure 3**).

#### Flow Cytometry

The dESCs, as well as DF (see RT-qPCR section), are resuspended in 1:1000 viability dye (Invitrogen eFluor™ 450) solution in PBS for 10 minutes. The viability dye is used to distinguish live from dead cells based on their membrane integrity. Cells with a damaged membrane uptake the dye, which cannot pass through the intact membrane of healthy cells.

The supernatant is then removed via centrifugation (480g, 5 minutes) and replaced with a 1:50 primary antibody solution (either EMA-1, AB_531885 by DSHB or SSEA-1, AB_528475 by DSHB) in a 10% FBS-PBS solution. EMA-1 recognizes cell surface glycolipids in mouse embryonic stem cells and chicken blastodermal cells. SSEA-1, a carbohydrate antigen on glycolipids, is strongly expressed by the murine inner cell mass of the blastocyst (ICM) and murine embryonic stem cells ^37^.

After 2 hours of incubation, cells go through three centrifugation-wash cycles (480g, 5 minutes) with 10%FBS-PBS solution, after which they are resuspended in a 1:400 solution of FITC-conjugated secondary antibody. After 30 minutes of incubation, two more centrifugation-wash cycles (480g, 5 minutes) are performed, then the cells are analyzed by the Mitelnyi MACSQuant® Analyzer 16 Flow Cytometer.

Cells are first selected for size to exclude aggregates, debris and cell doublets. Then, the percentage of cells that are both not marked by FV450 (i.e., viable cells) is quantified and marked by each of the primary antibodies (via fluorescence detection of the common secondary antibody). Resulting percentages are then plotted and can be found in **Figure 4** of the main text. Normality is tested via Shapiro-Wilk test, then an unpaired t-test is performed to assess difference between the two groups.

### Biomass composition

To evaluate the nutritional profile of the duck ESC biomass, five independent production batches (*n*=5) were harvested and subjected to standardized, food-grade analytical testing. The assessment utilized validated methodologies, including AOAC 950.46 for moisture content, AOAC 981.10 for total protein, AOAC 960.39 for crude fat, and AOAC 920.153 for ash. Additionally, the amino acids profile was characterized following ISO 13903:2005. The carbohydrate amount was calculated by subtracting moisture, crude fat, total protein, and ash from 100 %. Energy, expressed in kcal was calculated based on calorie values for 1 g of protein (4 kcal), carbohydrates (4 kcal), and fat (9 kcal). The obtained data were compared to that of conventional raw duck meat and goose liver deposited in the USDA food database.

### Proteomics

Proteomics was performed following an optimized bottom-up LC-MS workflow^38^. Proteins were extracted, reduced, alkylated, and digested using the EasyPep™ kit. LC-MS analysis used a Vanquish Neo™ UHPLC coupled to an Orbitrap Exploris™ 480 MS, with peptide separation on a PepMap Neo C18 column at 50°C. Mobile phases consisted of water (A) and 80% ACN (B), both with 0.1% FA, at 300 nL/min. The 64-minute gradient ranged from 0–95% B with a flow rate ramp to 450 nL/min. Analysis was performed in DDA mode: MS1 full scans (R=120K) and ddMS2 scans (R=15K, cycle time=2s), with HCD at 30% and a 45s dynamic exclusion. Data was processed in ProteomeDiscoverer™ against the *Anas platyrhynchos* UniProtKB database (https://www.uniprot.org) using a 10 ppm precursor and 0.02 Da fragment tolerance. Trypsin digestion allowed two missed cleavages; FDR was set to 1%. Proteins with at least one unique peptide were used for quantification. For the statistical analysis, one-way ANOVA evaluated group differences, with significance at *p* < 0.05. Results are reported as mean ± standard deviation.

## Acknowledgements

We thank the whole team at PARIMA for fruitful discussion and providing supporting data for the results presented in this paper.

---

**Figure S1.**
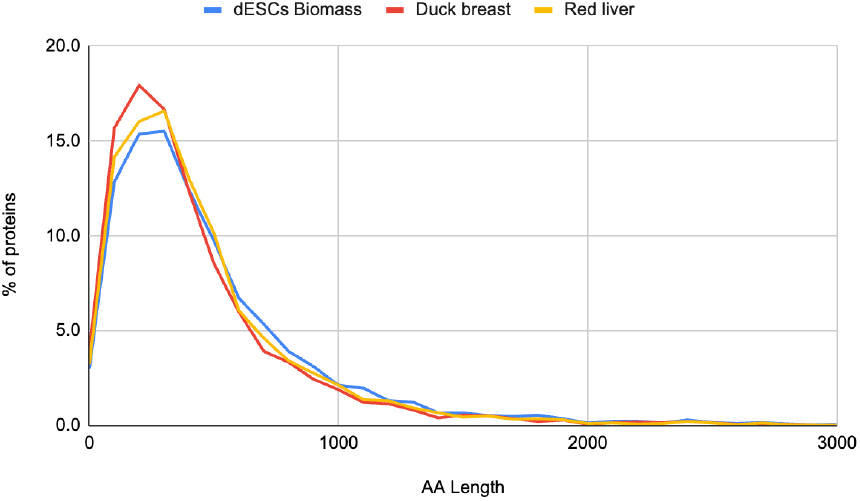
Protein size distribution of dESCs biomass, duck breast, and liver protein_s_ (in terms of amino acid length). Proteins levels are expressed in percentage of the total count

**Figure S2.**
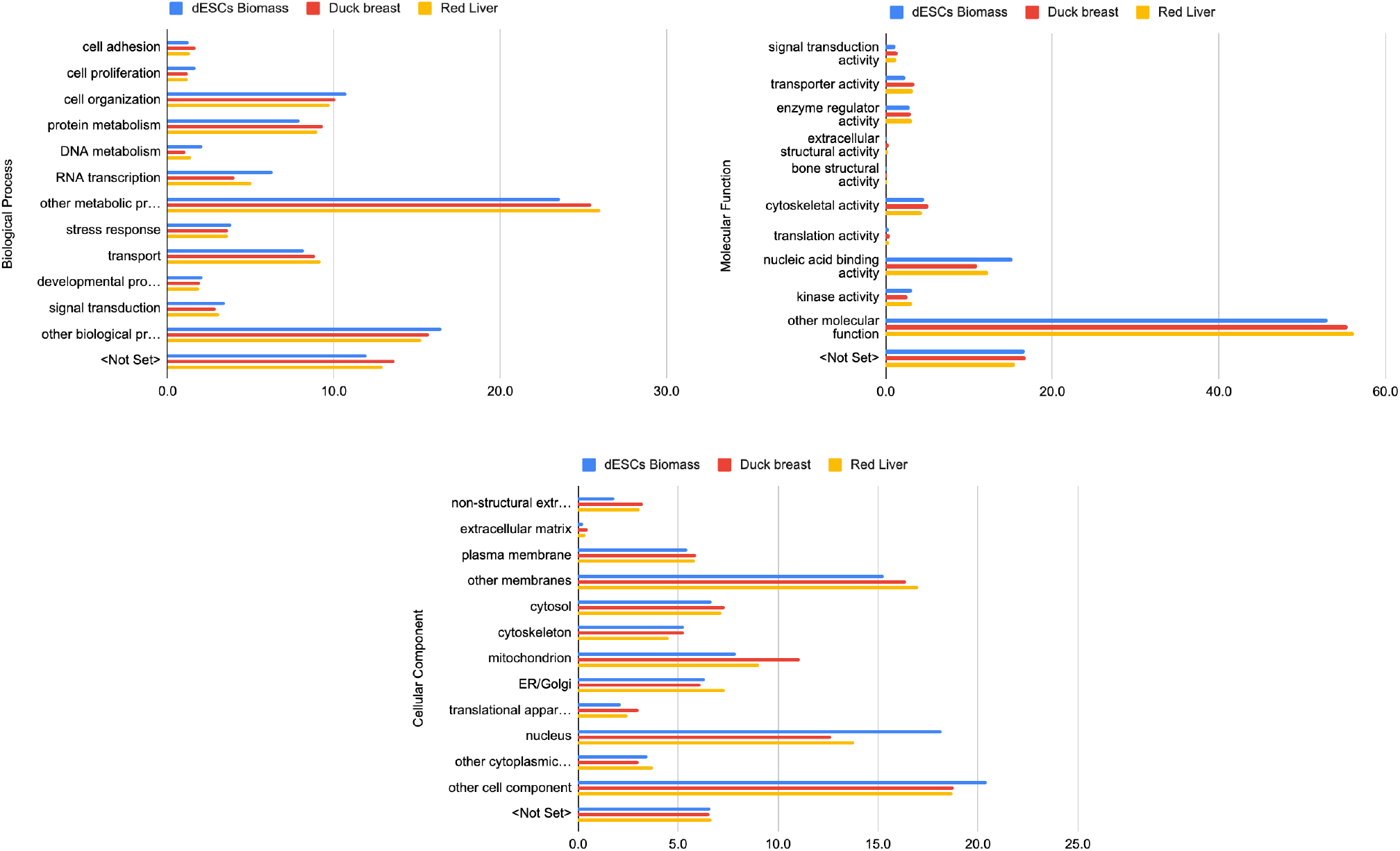
Comparison of the protein profile of different sample groups – dESCs biomass, duck breast, and Duck liver – through Gene Ontology (GO). GO analysis of the identified proteins is expressed in percentage of the total count. The three categories of GO are represented: Cellular Component (CC), Biological Process (BP), and Molecular Function (MF)

## References

1. Lin, Y.-C. et al. Cultivated meat for sustainable food security and environmental resilience Future Foods 13, 100984 (2026).

2. Gu, H. et al. Scaling cultured meat: challenges and solutions for affordable mass production. Compr. Rev. Food Sci. Food Saf. 24, e70221 (2025).

3. Muller, Q. & Matsusaki, M. Comparative analysis of cultivated meat cell and cell type usage across species: functional roles and engineering potential. Trends Food Sci. Technol 169, 105524 (2026).

4. Lee, M. et al. Cultured meat with enriched organoleptic properties by regulating cell differentiation. Nat. Commun. 15, 77 (2024).

5. Bakhsh, A. et al. Cell-based meat safety and regulatory approaches: a comprehensive review. Food Sci. Anim. Resour. 45, 145–164 (2025).

6. Pain, B et al. Long-term *in vitro* culture and characterisation of avian embryonic stem cells with multiple morphogenetic potentialities. Development 122, 2339–2348 (1996).

7. Jara, T. C. et al. Stem cell-based strategies and challenges for production of cultivated meat. Nat. Food 4, 841–853 (2023).

8. Chen, X. et al. Derivation of embryonic stem cells across avian species. Nat. Biotechnol. (2025). doi:10.1038/s41587-025-02833-3.

9. Mathieu, T. et al. Integrative multi-omics modeling for cultivated meat production, quality, and safety. Trends Food Sci. Technol. 166, 105364 (2025).

10. Ng, W. L. & Tan, J. S. Machine learning in cultivated meat: enhancing sustainability, efficiency, quality, and scalability across the production pipeline. Food Bioprocess Technol. 18, 5988–6009 (2025).

11. Amirvaresi, A., Shahsavari, A. & Ovissipour, R. Integrated multi-omics characterization of CDK4/hTERT-immortalized ovine satellite cells for cultivated meat applications. Preprint at bioRxiv 10.1101/2025.10.23.684220 (2025).

12. Bennie, R. Z. et al. A risk-based approach can guide safe cell line development and cell banking for scaled-up cultivated meat production. Nat. Food 6, 25–30 (2025).

13. International Conference Harmonisation of Technica Requirements for Registration of Pharmaceuticals for Human Use. ICH Q5D: Derivation and Characterisation of Cell Substrates Used for Production of Biotechnological/Biological Products (ICH, 1997).

14. Hebert, P. D N., Cywinska, A., Ball, S. L. & deWaard, J. R. Biological identifications through DNA barcodes. Proc. R. Soc. Lond. B Biol. Sci. 270. 313–321 (2003).

15. Messmer, T. et al. A serum-free media formulation for cultured meat production supports bovine satellite cell differentiation in the absence of serum starvation. Nat. Food 3, 74–85 (2022).

16. Pasitka, L. et al. Spontaneous immortalization of chicken fibroblasts generates stable, high-yield cell lines for serum-free production of cultured meat. Nat. Food 4, 35–50 (2023).

17. Qanbari, S. et al. Genetics of adaptation in modern chicken. PLoS Genet 15, e1007989 (2019).

18. Samuels, D. C. et al. Heterozygosity ratio, a robust global genomic measure of autozygosity and its association with height and disease risk. Genetics 204, 893–904 (2016).

19. Hansen, J., Jain, A. R., Nenov, P., Robinson, P. N. & Iyengar, R. From transcriptomics to digital twins of organ function. Front. Cell Dev. Biol. 12, 1240384 (2024).

20. Jiang, Q. et al. METTL14 regulates proliferation and differentiation of duck myoblasts through targeting MiR-133b. PLoS ONE 20, e0320659 (2025).

21. Lavial, F. et al. The Oct4 homologue PouV and Nanog regulate pluripotency in chicken embryonic stem cells. Development 134, 3549–3563 (2007).

22. Heurtier, V. et al. The molecular network of NANOG in pluripotency and reprogramming. Nat. Commun. 10, 5735 (2019).

23. Nishitani, E., Li, C., Lee, J., Hotta, H., Katayama, Y., Yamaguchi, M. & Kinoshita, T. Pou5f3.2-induced proliferative state of embryonic cells during gastrulation of Xenopus laevis embryo. Dev. Growth Differ. 57, 591–600 (2015).

24. Flores, I., Benetti, R. & Blasco, M. A. Telomerase regulation and stem cell behaviour. Curr. Opin. Cell Biol. 18, 254–260 (2006).

25. Ehrlich, M. Expression of various genes is controlled by DNA methylation during mammalian development. J. Cell. Biochem. 88, 899–910 (2003).

26. Jean, C. et al. Transcriptome analysis of chicken ES, blastodermal and germ cells reveals that chick ES cells are equivalent to mouse ES cells rather than EpiSC. Stem Cell Res 11 586–599 (2013).

27. Pain, B., Kress, C. & Rival-Gervier, S. Pluripotency in avian species. Int. J. Dev Biol 62, 245–255 (2018).

28. Semrau, S. et al. Dynamics of lineage commitment revealed by single-cell transcriptomics of differentiating embryonic stem cells. Nat. Commun. 8, 1096 (2017).

29. Klopfenstein, D. V. et al. GOATOOLS: a Python library for Gene Ontology analyses. Sci. Rep. 8, 10872 (2018).

30. Subramanian, A. et al. Gene set enrichment analysis: knowledge-based approach for interpreting genome-wide expression profiles. Proc. Natl Acad. Sci. USA 102, 15545–15550 (2005).

31. Evans, M. J. & Kaufman, M. H. Establishment in culture of pluripotential cells from mouse embryos. Nature 292, 154–156 (1981).

32. Ying, Q. L. et al. The ground state of embryonic stem cell self-renewal. Nature 453, 519–523 (2008).

33. Katsumoto, T., Mitsushima, A. & Kurimura, T. The role of the vimentin intermediate filaments in rat 3Y1 cells elucidated by immunoelectron microscopy. Biol. Cell 68, 139–146 (1990).

34. Etches, R. J., Clark, M. E., Verrinder Gibbins, A. & Cochran, M. Contributions to somatic and germline lineages of chimeric chickens following injection of somatic cells into embryos. Biol. Reprod. 56, 128–133 (1997).

35. Horiuchi, H. et al. Chicken leukemia inhibitory factor maintains chicken embryonic stem cells in the undifferentiated state. J. Biol. Chem. 281, 24280–24288 (2006).

36. Selliah, N. et al. Flow cytometry method validation protocols. Curr. Protoc. Cytom. 87, e53 (2018).

37. Henderson, J. K. et al. Preimplantation human embryos and embryonic stem cells show comparable expression of stage-specific embryonic antigens. Stem Cells 20, 329–337 (2002).

38. Palma, J., Colchero Leblanc, C., Kusters, R. & Kamgang Nzekoue, A. F. Proteomics for cultivated meat: the importance of analytical standardization. Preprint at bioRxiv 10.64898/2026.03.23.713501 (2026).

39. Fang, Y. et al. Comparative transcriptomic analysis of embryonic stem cells across mammalian species. iScience 29(1) (2026).

40. Tang, J. et al. Dietary riboflavin supplementation improves meat quality, antioxidant capacity, fatty acid composition, lipidomic, volatilomic, and proteomic profiles of breast muscle in Pekin ducks. Food Chem. X 19, 100799 (2023).

41. Yang, S. et al. Proteomic analysis of liver tissues in chicken embryo at Day 16 and Day 20 reveals antioxidant mechanisms. J. Proteomics 243 104258 (2021).

42. Ham, J. H., Lee, Y. J., Lee, I. & Kim, H. Y. Allergenicity in cultured meat: assessment and strategic management. Crit. Rev. Food Sci. Nutr. 65, 8246–8258 (2025).

43. Deng, M. et al. A review on allergens, allergenicity assessment methods, and specific allergenicity assessment strategies for novel foods. J. Agric. Food Chem. 73, 27177–27188 (2025).

44. Wu, W., Fu, Y., Therkildsen, M., Li, X. M. & Dai, R. T. Molecular understanding of meat quality through application of proteomics. Food Rev Int 31, 13–28 (2015).

45. Fraeye, I., Kratka, M., Vandenburgh, H. & Thorrez, L. Sensorial and nutritional aspects of cultured meat in comparison to traditional meat: much to be inferred. Front. Nutr. 7, 35 (2020).

46. Broucke, K., Van Pamel, E., Van Coillie, E., Herman, L. & Van Royen, G. Cultured meat and challenges ahead: a review on nutritional, technofunctional and sensorial properties, safety and legislation. Meat Sci. 195, 109006 (2023).

47. Cassotta, M. et al. Human-based new approach methodologies to accelerate advances in nutrition research. Food Front. 5, 1031–1062 (2024).

48. Levine, S. L. et al. Challenges and opportunities in the development and adoption of new approach methods (NAMs). J. Agric. Food Chem. 73, 7519–7521 (2025).

49. Zandonadi, R. P., Ramos, M C., Elias, F. T. S. & Guimarães, N. S. Globa insights into cultured meat uncovering production processes, potential hazards, regulatory frameworks, and key challenges — a scoping review. Foods 14, 129 (2025).

50. Sims, D., Sudbery, I., Ilott, N. E., Heger, A. & Ponting, C. P. Sequencing depth and coverage: key considerations in genomic analyses. Nat. Rev. Genet. 15, 121–132 (2014).

